# Transcriptomic analysis during fruit development of the oil palm revealed specific isozymes related to starch metabolism that control oil yield

**DOI:** 10.1101/2023.04.14.536940

**Authors:** Ardha Apriyanto, Julia Compart, Joerg Fettke

## Abstract

The oil palm (*Elaeis guineensis* Jacq.) produces a large amount of oil from the fruit. A recent study has shown that starch metabolism is essential for oil synthesis in fruit-producing species. Therefore, we detected gene expression changes related to starch metabolism genes throughout the maturity stages of oil palm fruit with different oil yields. Gene expression profiles were examined with three different oil yields (low, medium, and high) at six fruit development phases (4, 8, 12, 16, 20, and 22 weeks after pollination). Using RNA-seq analysis, we successfully identified and analyzed differentially expressed genes in oil palm mesocarps during development. The results showed that the transcriptome profile for each developmental phase was unique. Additionally, we found that starch synthesis and degradation occurred during fruit development and influenced oil production. Sucrose flux to the mesocarp tissue, rapid starch turnover, and high glycolytic activity have been identified as critical factors for oil production in oil palms. For starch metabolism and the glycolytic pathway, we identified specific enzyme isoforms (isozymes) that may control the oil production. This study provides valuable information for creating new high-oil-yielding palm varieties via breeding programs or genome editing approaches.

## Introduction

The oil palm (*Elaeis guineensis Jacq.*) produces the highest yield per hectare of land compared to other oil crops (Barcelos et al., 2015; Meijaard et al., 2020). Consequently, it has become the most important commercial oil crop, particularly in Indonesia and Malaysia. Today, palm oil is used in a wide range of common products and is used on many industrial scales (food and non-food sectors). Palm oil accounts for more than 40% of the global demand for vegetable oil, and its demand is expected to increase significantly in the future (Meijaard et al., 2020). It is predicted that 93–156 Mt of palm oil will be required by 2050 (Murphy et al., 2021; Pirker et al., 2016).

To fulfill this demand, palm oil production must be enhanced. However, increasing the palm oil production is not a simple task because oil yield is a genetically complex trait involving many genes, especially in the case of perennial oleaginous crops, such as oil palm. The long breeding cycle of this plant, which usually takes approximately 20 years, limits the development of new varieties (John Martin et al., 2022). Furthermore, numerous challenges are likely to appear in the future, including emerging threats from climate change, pests, and diseases, which will diminish palm oil production (Murphy et al., 2021). Therefore, new technologies must improve the oil production of this plant.

Palm oil is produced in fruit mesocarp tissue primarily by triacylglycerols (TAG). The mesocarp contains up to 90 % dry weight of oil, which is one of the highest rates of oil accumulation in the plant tissues (Bourgis et al., 2011; Tranbarger et al., 2011). Therefore, understanding the mechanism of oil deposition in palm fruit mesocarps is an exciting research topic.

Several recent studies have elaborated on the mechanisms of oil biosynthesis during fruit development (Dussert et al., 2013; Teh et al., 2014; Tranbarger et al., 2011). Nonetheless, most of this research has focused on the lipid biosynthesis pathway, with little attention paid to its interactions with other metabolic processes. However, a link between starch metabolism and oil production in the palm mesocarp has been previously reported (Bourgis et al., 2011; Guerin et al., 2016). Further, there was an indication of starch deposition at the end of fruit maturation (Bourgis et al., 2011). Starch metabolism and oil biosynthesis genes are co-expressed during oil deposition, which suggests that starch metabolism is crucial for oil synthesis (Guerin et al., 2016). Our recent findings showed that starch parameters, such as total starch content, starch granule size, and chain length distribution, are correlated with oil yield (Apriyanto et al., 2022b). Additionally, starch-related hydrolytic activity during fruit development is strongly associated with oil yield (Apriyanto et al., 2022b).

In higher plants, starch is the principal storage carbohydrate, composed of two glucose polymers, amylose, and amylopectin, which form complex semi-crystalline granules inside the plastids (Apriyanto et al., 2022a). Starch is also synthesized within the mesocarp tissue, although its detailed function remains obscure. Furthermore, the exact starch synthesis and degradation routes used inside the mesocarp tissue of the oil palm remain unclear. However, similar to other organs and tissues, starch metabolism in the palm fruit mesocarp involves many enzymes and proteins. The predicted starch metabolism is shown in Figure 1. The proposed pathway was adapted from potato (*Solanum tuberosum*) (van Harsselaar et al., 2017), banana (*Musa accuminata*) (Cordenunsi-Lysenko et al., 2019; Kuang et al., 2021), and Arabidopsis (*Arabidopsis thaliana*) (Sonnewald and Kossmann, 2013).

**Figure 1.**
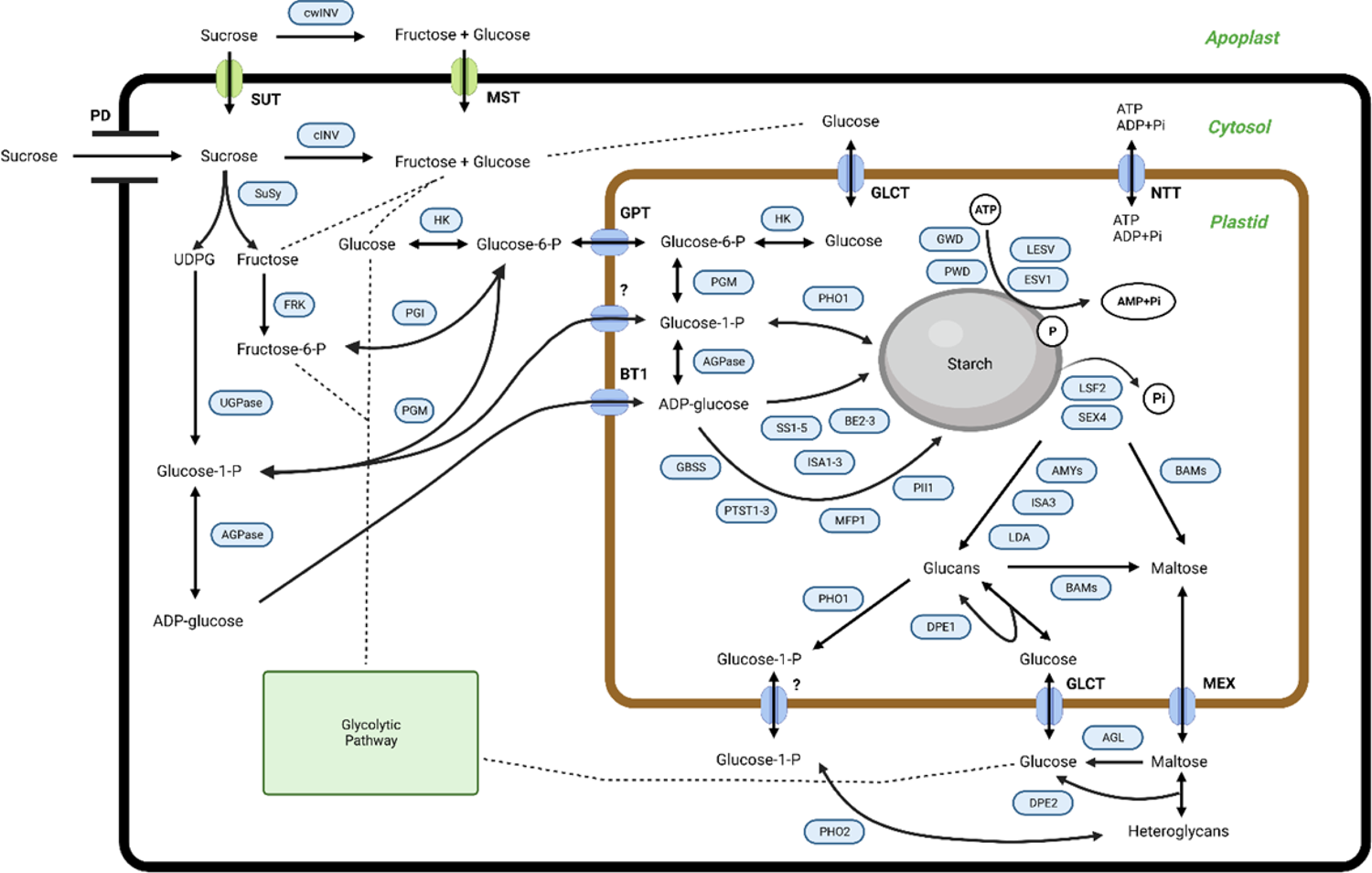
The proposed pathway for starch metabolism in the mesocarp tissue of oil palms. SUT, sucrose transporter; cwINV, cell wall invertase; cINV, soluble invertase; SuSy, sucrose Synthase; HK, hexokinase; FRK, fructokinase; PGI, phosphoglucoisomerase; PGM, phosphoglucomutase; UGPase, UDP-glucose pyrophosphorylase; AGPase, ADP-glucose pyrophosphorylase; SS, starch synthase; GBSS, granule-bound starch synthase; BE, starch branching enzyme; GWD, glucan, water dikinase; PWD, phosphoglucan, water dikinase; ESV1, early starvation 1; LESV, like early starvation; BAM, beta-amylase; AMY, alpha-amylase; SEX4, starch excess 4; LSF2, like starch-excess 2; LDA, limit dextrinases; DPE, disproportionating enzyme; MFP, MAR-binding filament-like protein 1; PII, protein involved in starch initiation; PTST, protein targeting to starch; BE, starch branching enzyme; ISA, isoamylase; PHO, alpha-glucan phosphorylase; AGL: alpha-glucosidase; GPT, glucose 6-phosphate/phosphate translocator; BT1, ADP glucose transporter; NTT, nucleotide translocator; GLCT, glucose transporter; MEX, maltose transporter; PD, plasmodesmata. For further information regarding starch metabolism in plants, see (Apriyanto et al., 2022a) and references therein.

To improve our understanding of starch metabolism and its link to oil production in palm fruits, we used six developmental stages of oil palm fruit development with different oil yields. We applied an RNA sequencing (RNA-seq) approach using next-generation sequencing (NGS) platforms to analyze transcriptome profiles during fruit development. RNA-seq is frequently used to elucidate transcript structures, variations, and gene expression levels owing to its high accuracy, whole-genome coverage, and extensive detection range. Consequently, many successful transcriptome profiling studies have used this RNA-seq approach during fruit development in several plants. These include tomatoes (Shinozaki et al., 2018) and bananas (Kuang et al., 2021).

Overall, we aimed to (1) detect changes in gene expression during the development of oil palm fruits and (2) reveal the relationship between the starch metabolic pathway and oil production. The results of this study will advance our understanding of the relationship between starch metabolism and oil yield in oil palms and will provide breeding targets for palms with increased oil yield.

## Results

### Identification of genes encoding enzymes and proteins related to starch metabolism in oil palm

A homology search was conducted to identify starch metabolism genes in the oil palm. Based on sequence similarity, we found almost all homologous genes in comparison with *Arabidopsis thaliana* (Table S1). Genes coding for starch metabolism-related enzymes and proteins are distributed over all sixteen oil palm chromosomes (Figure 2). We found a cluster of synthesis genes of starch metabolism on chromosome 2, but most encoding genes are randomly distributed across different chromosomes. This situation is similar to that observed in *Solanum tuberosum* (van Harsselaar et al., 2017) and *Arabidopsis thaliana* (Sonnewald and Kossmann, 2013). Here, we successfully identified 113 loci encoding enzymes and proteins related to starch metabolism in the oil palm (Table 1), which have not been previously reported.

**Figure 2.**
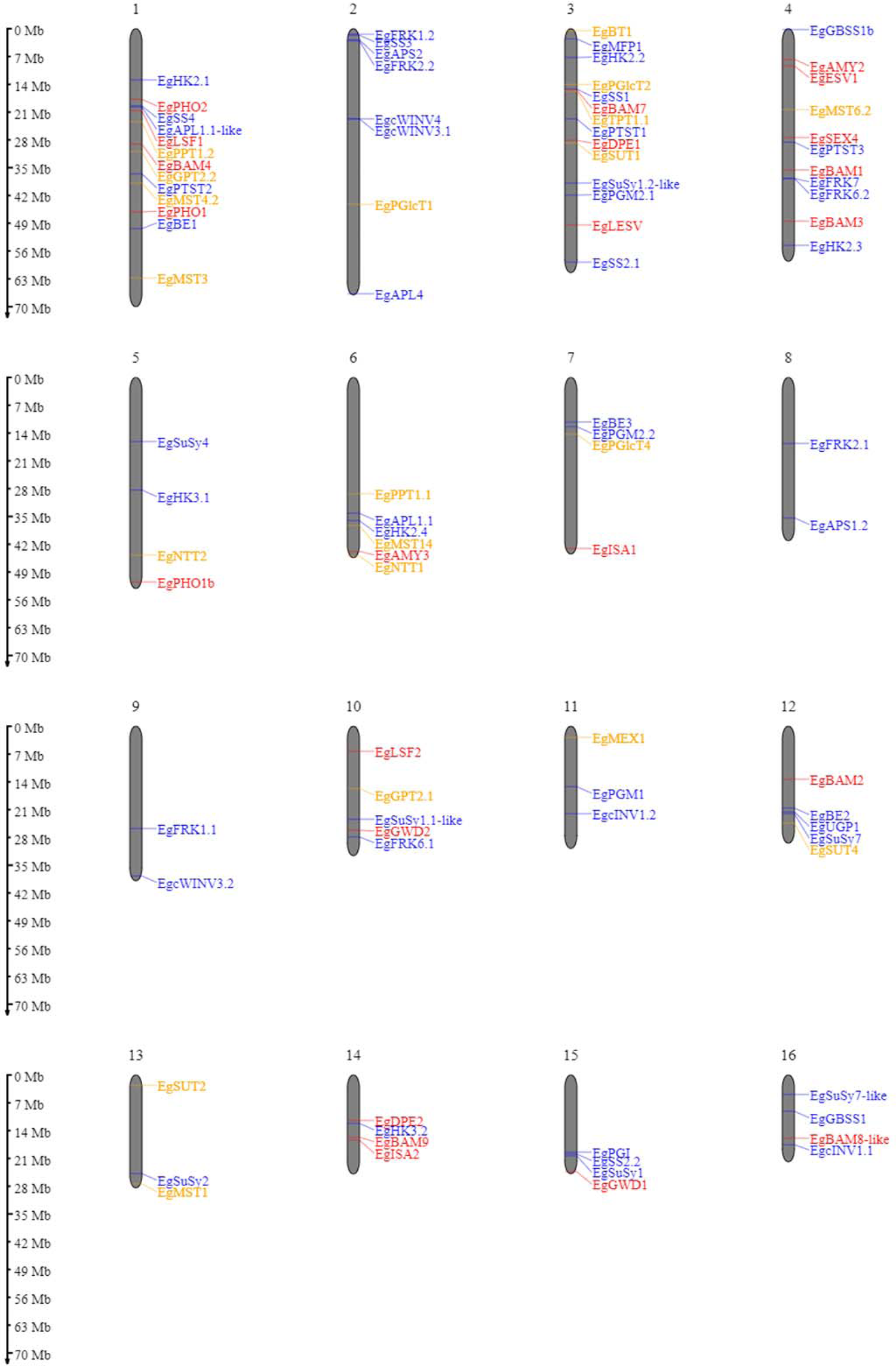
Ideogram of the physical positions of starch metabolism-related genes in the oil palm genome. The relative map positions of 101 genes encoding starch metabolism-related genes are shown for individual pseudomolecules depicting chromosomes 1–16. Blue, red, and yellow letters depict the synthesis, degradation, and transporter pathways, respectively. Genes from the unscaffolded group were not displayed.

Enzymes involved in starch metabolism often belong to gene families that encode for multiple isoforms. We identified 113 loci that encode starch metabolism-related enzymes and proteins. However, as shown in Table 1, several enzyme isoforms known in other species were not discovered in the oil palm. Two examples should be mentioned here: SS and BAM gene families. In the oil palm, we could not find the starch synthase 6 isoform, as reported in potato (van Harsselaar et al., 2017). Furthermore, no BAM5, BAM6, or BAM10 was identified in the oil palm. However, for potato, BAM 6 has been described, but not BAM5 and BAM8 isoforms (van Harsselaar et al., 2017). The BAM10 isoform is also absent in *Arabidopsis thaliana* (Thalmann et al., 2019). This situation in oil palm differs from that of bananas and potato, which might be due to polyploidy, as this will influence the variation in isoform numbers for the SS, AMY, and BAM gene families (Jourda et al., 2016; Wang et al., 2022).

Another remarkable finding was that only four isoforms of the sucrose synthase gene family were detected in the oil palm. These are SuSy1, SuSy2, SuSy4, and SuSy7. In contrast, six isoforms (SuSy1-7, except SuSy6) were identified in the potato genome (van Harsselaar et al., 2017).

Other gene families in the starch-related metabolic pathway of palm oil were found to be similar to those in Arabidopsis, and some isoforms were duplicated in palm oil, as shown in Table 1. In principle, starch metabolism proceeds similarly, although there are deviations in the number of isoforms.

### Transcriptomic profiles segregated by fruit ripening and oil yield

The classification of the high-, medium-, and low-yield groups used in this study was based on the oil yield data depicted in Figure 3a,b. The low-yielding fruits had on average 26% oil to bunch (%OB) and 78% oil to dry mesocarp (%ODM) values, medium-yielding fruits had 32% OB and 82% ODM, and high-yielding fruits had 38% OB and 85% ODM, respectively. Each group had significantly different oil-to-dry mesocarps (% ODM) and oil-to-bunch (% OB) values.

**Figure 3.**
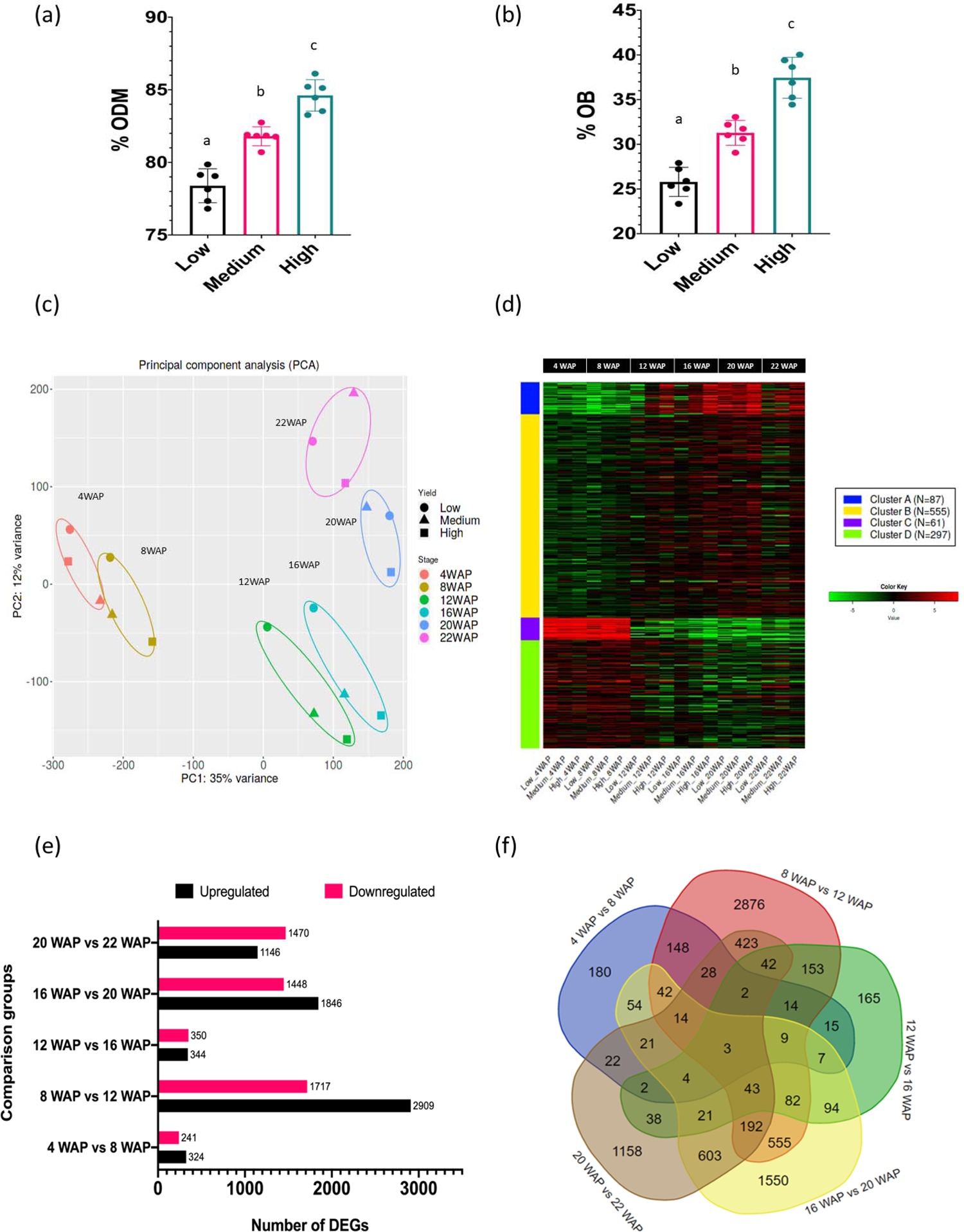
Oil yield and gene expression profiles of oil palm fruit mesocarps during fruit ripening. (a) The percentages of oil to dry mesocarp (% ODM) and (b) oil to bunch (% OB) of the oil palm yield are displayed. For statistical analysis, each group (n = 6) was tested using one-way ANOVA with Tukey’s post-hoc analysis; letters indicate a significant difference between matching groups (p <0.001). (c) Principal component analysis (PCA) of transcriptome data during fruit development. (d) Gene clusters during fruit development were identified using k-means clustering analysis. The heatmap scale shows the normalized expression level. (e) Number of DEGs identified in the particular comparison groups. (f) Venn diagram of DEGs in five comparison groups.

To understand transcriptome alterations in oil palm fruit at different stages and oil yields, an RNA-seq approach was performed. From six stages of fruit development, with three different oil yields, 18 cDNA libraries were constructed and further sequenced. The sequencing results generated 6.1–8.2 Gbps data per library (Table S2), and clean reads were mapped onto the oil palm reference genome sequence (Singh et al., 2013). Principal component analysis (PCA) was used to visualize the transcriptomic profiles of each fruit ripening stage (Figure 3c) and showed that the fruit developmental stages were clustered according to their WAP, indicating that the transcriptomic profile of each developmental stage was unique, as expected. Furthermore, this indicated that shifts in gene expression occurred over time, allowing us to distinguish the transition between the early and late periods of development and ripening. Such specific transcriptomic profiles during the fruit ripening stage have also been observed in other fruit-producing species, such as tomatoes and peppers (Osorio et al., 2012). Interestingly, with increased ripening, the metabolism of the different oil-yielding fruits was more strongly separated (Figure 3c; 16, 20, and 22 WAP). This clearly demonstrated that specific genes were differentially expressed between the oil yield groups.

As previously mentioned, some genes were likely co-expressed during fruit development. Thus, we could identify at least four gene clusters that appeared during fruit development based on K-means clustering analysis (Figure 3d). Gene clusters A, B, C, and D consisted of 87, 555, 61, and 297 genes, respectively. This analysis is important for identifying groups of genes with similar expression patterns that may be controlled by key transcription factors. The coordinated process of gene expression alterations during fruit development has also been reported in tomatoes (Shinozaki et al., 2018) and bananas (Kuang et al., 2021).

One of our objectives was to identify the genes that play key roles in each fruit maturation stage. Therefore, a differentially expressed gene (DEG) analysis of the oil palm mesocarp between close fruit development stages was performed. The comparison of the six developmental stages of oil palm fruit was divided into five groups. The DEGs among the fruit development stages are shown in Figure 3e. The DEG results showed that there were 565, 4626, 694, 3294, and 2616 DEGs in 4WAP vs. 8WAP, 8WAP vs. 12WAP, 12WAP vs. 16WAP, 16WAP vs. 20WAP, and 20WAP vs. 22WAP, respectively (Figure 3e). Furthermore, the 4WAP vs. 8WAP group contained the least upregulated (324) and downregulated (241) DEGs, whereas the 8WAP vs. 12WAP group contained the most upregulated (2909) and downregulated (1717) DEGs. The list of DEGs is shown in Table S4. We also found that some DEGs were specific to each group comparison, that is, 180, 2876, 165, 1550, and 1158 genes in 4WAP versus 8WAP, 8WAP versus 12WAP, 12WAP versus 16WAP, 16WAP versus 20WAP, and 20WAP versus 22WAP, respectively. Three genes were shared between the five pairwise comparisons (Figure 3f). This means that these genes were always significantly upregulated or downregulated during fruit development. These genes were early nodulin-93 (LOC105035564), inorganic pyrophosphatase 1 (LOC105040479), and the BTB/POZ domain-containing protein At1g63850 (LOC105054221). The reason for this significant fluctuation during fruit ripening and maturation is still unknown, as these genes have not been well characterized in plants.

To identify the functions of key genes that influence fruit development, we mapped the DEGs to the KEGG database to identify the pathways enriched by DEGs. This revealed that specific pathways were significantly altered during fruit development (Figure. S1). For instance, the DNA replication pathway was significantly enriched from 4 to 12WAP (Figure S1a,b). Additionally, the fatty acid metabolism, fatty acid synthesis, and fatty acid elongation pathways were significantly enriched from 16 to 22 WAP (Figure S1c,d,e). This result also confirms previous studies suggesting that cell division and expansion are most pronounced during the fruit growing phase (4 to 8 WAP), and during the fruit ripening phase (16 to 22 WAP) when oil accumulation occurs (Tranbarger et al., 2011).

Comparing 12-16 WAP showed that fatty acid biosynthesis, metabolism, pyruvate, glycolysis, citric acid cycle, and central metabolism were significantly enriched (Figure S1c). All of these pathways are important for oil production. Therefore, we found that the most critical alteration in gene expression in oil production occurred during this period, as this is the transition phase between the fruit growth and ripening phases.

### Gene expression of specific isozymes in the sucrose and starch metabolism pathways is associated with oil yield

Our previous study showed that starch and sucrose are biomarkers for palm oil production (Apriyanto et al., 2022b). Therefore, we focused on elaborating the gene expression profiles related to sucrose and starch metabolism and its closely related glycolysis pathway. The gene expression analysis related to starch metabolism during fruit development and with different yields is shown in Figure S2, which shows that the expression of genes related to this pathway fluctuates during fruit development. Interestingly, we found several essential genes involved in sucrose and starch metabolism whose expression patterns were positively correlated with oil yield, as shown in Figure 4. These genes included isoforms of sucrose transporter, fructokinase, hexokinase, phosphoglucoisomerase, ADP-glucose pyrophosphorylase large and small subunits, starch synthase, alpha-amylase, beta-amylase, and alpha-glucosidase (Figure 4). This indicates that the expression levels of these genes influence the oil yield.

**Figure 4.**
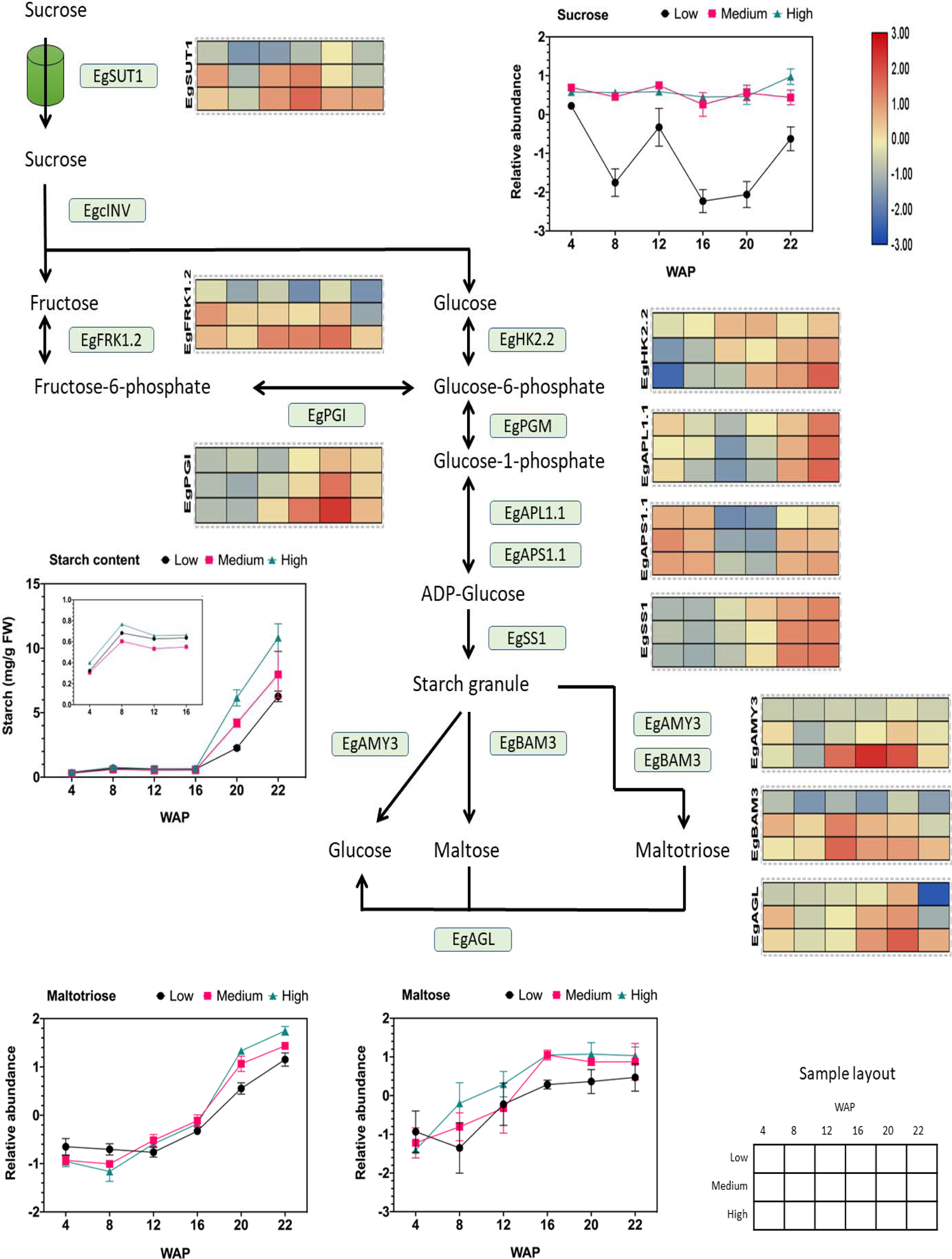
Gene expression profiles of sucrose and starch metabolic pathways in relation to starch and sugar abundance. The heat map shows the expression levels of genes encoding the corresponding genes, and the scale bar represents the normalized gene expression value. Only genes that were positively correlated with oil yield are shown. These genes are EgSUT1 (LOC105041710), EgPGI (LOC105060694), EgFRK1.2 (LOC105035371), EgHK2.2 (LOC105040520), EgAPL1.1 (LOC105047182), EgAPS1.1 (LOC105060488), EgSS1 (LOC105040918), EgAMY3 (LOC105047709), EgBAM3 (LOC105043800), and EgAGL (LOC105059813). The total starch, sucrose, maltotriose, and maltose contents during fruit development are depicted in the line graph.

We found that sucrose transporter 1 (SUT1) gene expression was consistently higher in high-yielding fruits than in low-yielding fruits during ripening and was correlated with sucrose abundance (Figure 4). We observed that the other SUT isoforms were also expressed during fruit development but were not correlated with oil yield (Figure S3a). Thus, the high influx of sucrose toward the mesocarp tissue, which is essential for oil production, is linked to the high expression of SUT1 (Figure 4). The main carbon source for lipid synthesis is supplied to oil palm fruit primarily as apoplastic sucrose (Bourgis et al., 2011). Therefore, the sucrose transporter is essential for the delivery of apoplastic sucrose to the mesocarp cells. Based on these results, SUT1 acts as a controller of sucrose flux into the mesocarp. A previous study also showed that overexpression of SUT1 in peas (*Pisum sativum*) leads to high sucrose loading in the sink part, thereby improving yield (Lu et al., 2020). However, this evidence also supports the idea that sucrose transporters are essential for enhancing fruit quality and crop yields (Aluko et al., 2021; Wen et al., 2022).

Additionally, it has been shown that starch parameters such as total starch content (Figure 4), starch granule size, and inner starch structure (Figure S4a,b) are correlated with oil yield (Apriyanto et al., 2022b). These data suggest that the total starch content is influenced by an increase in the expression of ADP-glucose pyrophosphorylase large and small subunits (APL and APS). ADP-glucose pyrophosphorylase is essential because it catalyzes the first committed step in starch synthesis. We found that the isoform APL1.1 and APS1.1 correlated with oil yield, but no other isoforms (Figure S3e). Similarly, overexpression of ADP-glucose pyrophosphorylase genes increased starch content in wheat (Kang et al., 2013).

As shown in Figure 4 and Figure S3f, the SS1 isoform was correlated with starch size and yield. Overexpression of lbSS1 in sweet potatoes increases the starch content and alter the granule size and structure of starch (Wang et al., 2017). Overall, we found that ADP-glucose pyrophosphorylase large 1.1 and small subunit 1.1 (APL1.1 and APS1.1), including starch synthase 1 (SS1), are important genes for initiating starch synthesis in oil palm mesocarps. To validate the transcriptomic results, starch synthase activity was evaluated by Native PAGE. The enzymatic activities of SS were shown in Figure S5. SS was proven to be active at 22 WAP. However, clear differences in the activity of soluble starch synthase at 22 WAP between different oil yields were detected (Figure S5).

In addition to starch synthesis, degradation pathways change during fruit development and yield. This was concluded from the correlation between increased hydrolytic activity and increased oil yield (Apriyanto et al., 2022b). As shown in Figure S3g and h, alpha-amylase, beta-amylase, and alpha-glucosidase gene expression fluctuated, except for the AMY3, BAM3, and AGL isoforms. The expression of these genes was correlated with the oil yield. Therefore, the increased abundance of maltose during fruit development (Figure 4) may have been caused by increased BAM3 expression. AMY3 and partially BAM3 may increase the abundance of maltotriose (see also Figure 4). Interestingly, earlier proteomic research revealed that alpha-amylase 3 (AMY3) abundance is significantly increased in high-yield oil palms (Loei et al., 2013). Alpha-glucosidase (AGL) also plays a role in producing glucose from maltose and maltotriose (Figure 4). This leads to a higher flux in the glucose pool.

### The oil yield is associated with the expression of specific isozymes in the glycolytic pathway

Glycolysis is also a crucial step in oil synthesis because it produces energy for cellular metabolism. Therefore, we examined the downstream pathway of starch metabolism, namely glycolysis. We found that almost all gene expression patterns inside the glycolytic pathway were positively correlated with oil yield (Figure 5). The list of genes involved in glycolysis is shown in Table S5, and gene expression analysis of this pathway is shown in Figure S6. Based on this, we also identified specific isoforms of the glycolysis pathway that are linked to oil yield. These include phosphofructokinase, fructose-1,6-bisphosphate aldolase, triose phosphate Isomerase, glyceraldehyde-3-phosphate dehydrogenase, phosphoglycerate kinase, phosphoglycerate mutase, enolase, and pyruvate kinase genes. The list of specific isoforms related to glycolysis that are correlated with oil yield is shown in Figure 6a.

**Figure 5.**
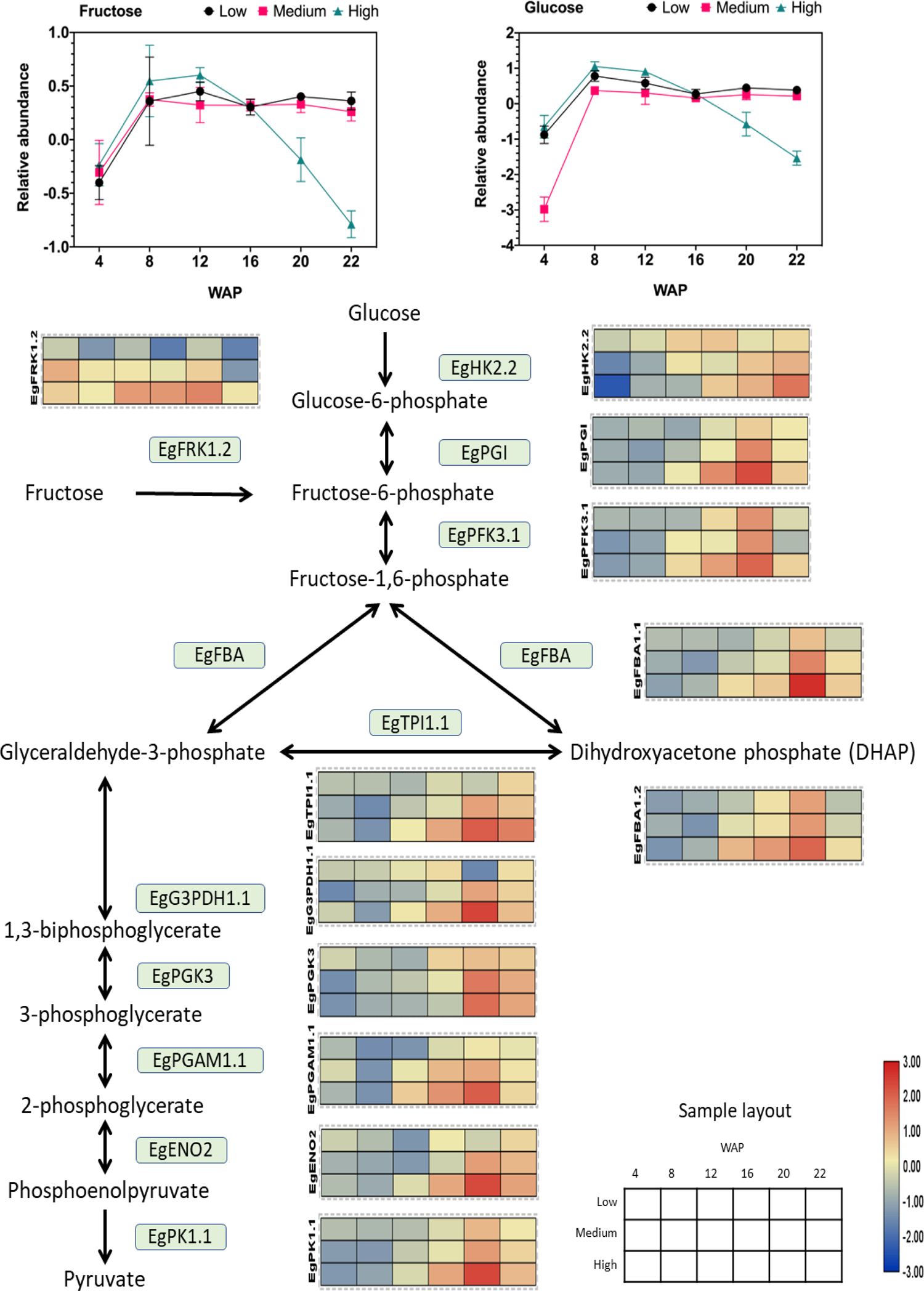
Gene expression profile of the glycolytic pathway in relation to sugar abundance. The heat map shows the expression levels of genes encoding the corresponding genes, and the scale bar represents the normalized gene expression value. Only genes that were positively correlated with oil yield are shown. These genes are EgFRK1.2 (LOC105035371), EgHK2.2 (LOC105040520), EgPGI (LOC105060694), EgPFK3.1 (LOC105040103), EgFBA1.1 (LOC105052993), EgFBA1.2 (LOC105059022), EgTPI1.1 (LOC105045658), EgG3PDH1.1 (LOC105034336), EgPGK3 (LOC105059872), EgPGAM1.1 (LOC105052340), EgENO2 (LOC105053561), and EgPK1.1 (LOC105038179). The abundances of fructose and glucose during fruit development are shown in the line graph.

**Figure 6.**
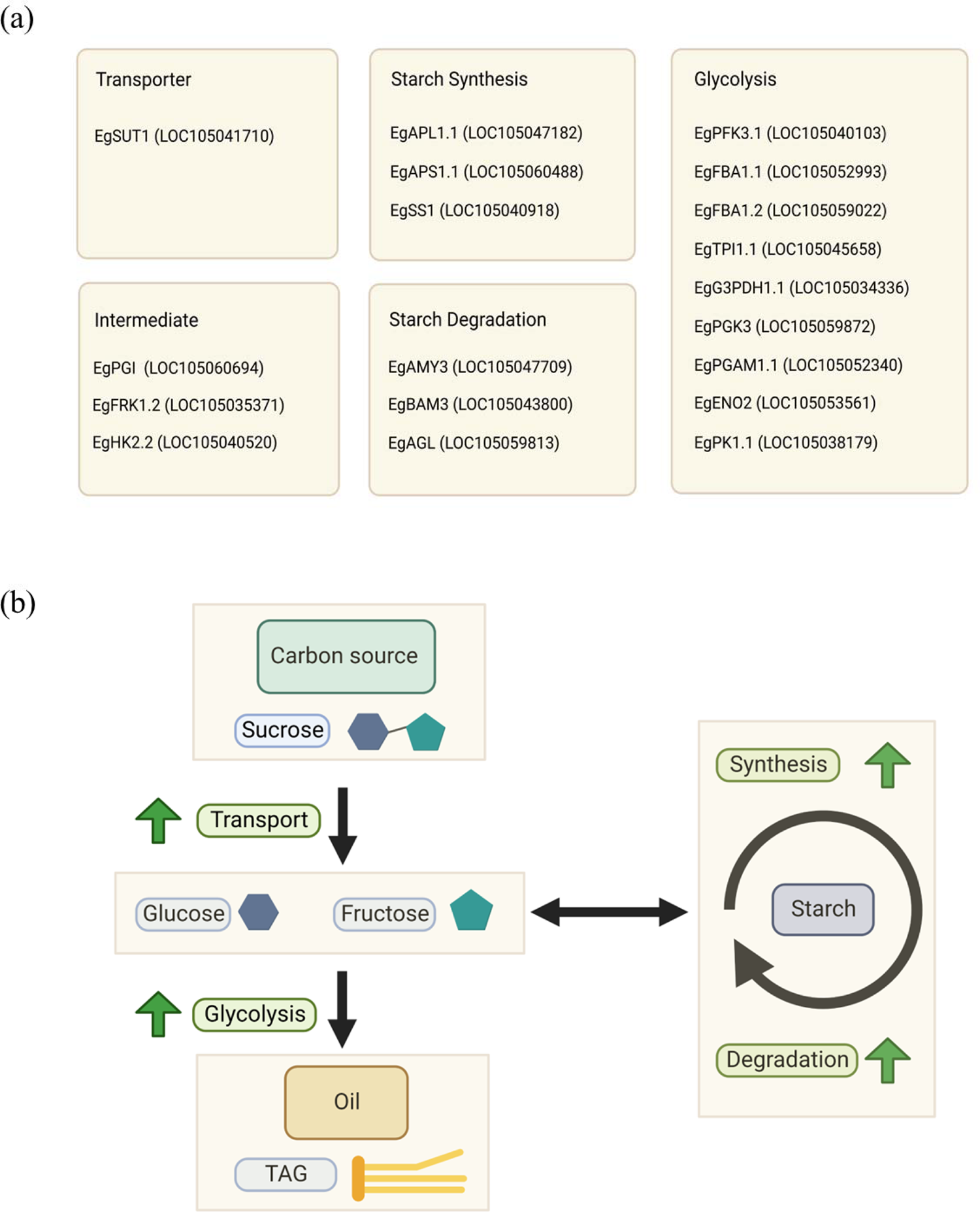
Key factors influencing the oil yield. (a) List of specific isoforms associated with the oil yield. (b) Schematic representation of the relationship between oil and starch metabolism in the high-yield group.

Interestingly, the fructose and glucose levels were depleted in the high-yielding group (Figure 5). Previous studies have shown that an increase in oil content in the avocado mesocarp during fruit growth is accompanied by a decrease in the concentration of reducing sugars (Kilaru et al., 2015). This result also corroborates with that of this study, which showed a decrease in the concentration of reducing sugars (such as glucose and fructose; Figure 5).

As shown in Figure 5, the expression of all the downstream genes was elevated in the high-yielding group. This indicates a high demand for both sugars for glycolysis in the high-oil-yielding group. Several previous studies have shown that high glycolytic activity is associated with high mesocarp oil content (Loei et al., 2013; Wong et al., 2017; Wong et al., 2014). Similar overexpression of oil palm fructose-1,6-bisphosphate aldolase (FBA) and glyceraldehyde-3-phosphate (G3PDH) resulted in increased lipid content in yeast (Ruzlan et al., 2017).

The RNA-seq results were verified by selecting several essential genes at the 22 WAP stage for qRT– PCR validation, which were positively correlated with the oil yield, as shown previously. The qPCR results showed that the expression patterns of these genes were consistent with the RNA-seq results (Figure S7).

## Discussion

### Starch synthesis and degradation happen during fruit ripening of oil palm

Currently, the prevailing opinion is that starch accumulates in either transient or long-term storage forms (Lloyd and Kossmann, 2015; Pfister and Zeeman, 2016). Transitory starch has a diurnal pattern: it is produced and accumulates directly from photosynthetic products during the day and is then degraded into sugars as an energy source for the following night; this is often seen in leaf organs (Zeeman et al., 2007). Storage starch, however, is generated and stored over time, as it is often found in perennating organs such as seeds, grains, embryos, and tubers (Lloyd and Kossmann, 2015). However, a third type of starch, known as ‘transitory-storage starch’, has been postulated (Luengwilai and Beckles, 2009a). It refers to starch that accumulates and degrades in the storage organs throughout development, particularly in fruits. (Luengwilai and Beckles, 2009b). Transitory-storage starch is a feature of many species, including economically valuable horticultural crops such as tomato, banana, apple, strawberry, nectarine, and kiwifruit. (Roch et al., 2020). Based on the data of this study, we found that the oil palm mesocarp tissue of the fruit also exhibits this type of ‘transitory-storage starch.’

Transitory-storage starch may be an evolutionary strategy for reproductive success that has unforeseen consequences for the postharvest industry (Dong and Beckles, 2019). First, increased fruit starch biosynthesis may enhance plant survival under stress conditions. Second, carbon storage as starch, rather than as sugars, minimizes osmotic disturbance in cells. Finally, from a postharvest perspective, starch in climacteric-ripening fruit may be a critical energy source for supporting biological processes and the synthesis of ‘quality-related’ metabolites that would reduce loss and waste (Yu et al., 2022).

### Oil palm fruit is categorized as climacteric fruit with unique starch metabolism pattern

A previous study found that oil palm mesocarp tissue exhibits the characteristics of climacteric fruit (Tranbarger et al., 2011). In relation to starch metabolism, many recent studies have shown that starch is a key component that may distinguish climacteric from non-climacteric fruits (Osorio et al., 2013; Chervin, 2020).

In most climacteric fruits, starch accumulates before the onset of ripening, and then starch is broken down into soluble sugars after the inception of ripening, whereas in non-climacteric fruits, the starch content drops very rapidly after anthesis, and they accumulate mainly soluble sugars throughout development and ripening. Interestingly, most climacteric fruits are transitory-storage starch type and most nonclimacteric fruits are sugar storer type (Yu et al., 2022).

Unlike typical climacteric fruits, in which the peak of starch accumulation occurs before the fruit ripening phase, the peak of starch accumulation in oil palms occurs during the ripening phase at 22 (Apriyanto et al., 2022b) or 23 WAP (Bourgis et al., 2011). This situation makes oil palm have unique characteristics as a climacteric fruit.

### Key factors to increase oil production in oil palm

Taken together, our data show that sucrose is used as a carbon source for lipid and starch biosynthesis, and subsequently, starch is used as a carbon source for further lipid synthesis. Similarly, labeled-metabolic flux analysis of tobacco leaves showed that lipids originate from starch (Chu et al., 2022). In the avocado mesocarp, which is rich in oil fruits similar to oil palms, starch is also a principal substrate for glycolysis, and transcripts for starch synthesis and degradation gene orthologs are abundant throughout mesocarp development (Pedreschi et al., 2019).

We found that the mechanism of carbon flux to produce oil in this study is similar to that of avocado, and probably the mechanism is conserved among crop-producing oil from the mesocarp tissue. Although the exact mechanism of carbon partitioning in oil palm mesocarp still requires further study, based on this evidence, we propose a relationship between oil and starch metabolism in oil palm, as depicted in Figure 6b.

In summary, high sucrose flux to the mesocarp tissue as a carbon source, rapid starch turnover, and high glycolytic activity are essential for higher oil production. We concluded that starch and sucrose metabolism, as well as glycolysis, in the mesocarp tissue are highly associated with oil production. We also found that specific isozymes in these pathways can drive and regulate the palm oil production. The specific isozymes were shown in Figure 6a.

This study will help understand starch metabolism in oil palm fruits and provide valuable resources for future genetic improvements in oil palms. The information obtained in this study is also suitable for modification of the physiological mechanism by specific treatment to increase the oil production of oil palms. Furthermore, the specific isozymes identified in this study could be used for target breeding or genome editing to create new oil palm varieties with enhanced oil yields.

## Methods

### Plant materials

A population of oil palm hybrid Deli × LaMe (DxL) tenera progenies from crosses of Deli dura females and LaMe pisifera males was used in this study. All progenies were planted in the Gunung Sejahtera Ibu Pertiwi Estate in Kalimantan Tengah, Indonesia, and were of the same age. Phenotypic observations, such as yield recording and bunch component analysis of each individual palm, were performed as described previously (Apriyanto et al., 2022b). Individual progeny trees with high, medium, and low yields were selected according to the evaluation after seven years of planting.

### Sample collections

Fruits from bunches were collected 4, 8, 12, 16, 20, and 22 weeks after pollination (WAP). Fruits were collected from each selected tree and randomly separated without bias from bunches during fruit development. Samples were frozen in liquid nitrogen and stored at −80°C. Prior to further analysis, the mesocarp tissue was ground into a fine powder in liquid nitrogen using a mortar and pestle.

### Starch parameter measurement

Starch parameters, such as total starch content, starch granule size distribution, and chain length distribution from the mesocarp tissue, were determined as described in our previous study (Apriyanto et al., 2022b).

### Metabolite profiling

The homogenized mesocarp tissue was used for metabolite profiling using gas chromatography-mass spectrometry (GC-MS). The measurement protocol and data analyzes were conducted as previously reported (Apriyanto et al., 2022b).

#### Identification of genes encoding starch metabolism-relevant enzymes and proteins

To identify the coding genes of enzymes and proteins involved in starch metabolism, *Arabidopsis thaliana* (Sonnewald and Kossmann, 2013), *Solanum tuberosum* (van Harsselaar et al., 2017), and *Musa acuminata* (Jourda et al., 2016) were used as the starting point for homology searches. All bioinformatics analyses, pairwise and multiple alignments, phylogenetic tree building, and assembly of DNA sequences were performed using Geneious Prime 2022.2.2 software (Kearse et al., 2012). The list of sequences (DNA, mRNA, and protein) from the three different organisms was compared with the oil palm genome sequence (GCF_ 00044 2705.1) using BLAST to identify the homologous sequences (Singh et al., 2013). A motif search was conducted using the MEME online tool (meme-suite.org), and motifs were compared between sequences within the same gene family (Bailey et al., 2015). The Kyoto Encyclopedia of Genes and Genomes (KEGG) database was used to verify the genes involved in metabolic pathways. The location of the identified genes was visualized in pseudochromosomes using the MG2C software (Chao et al., 2021).

#### RNA extraction, sequencing, and data analysis

Total RNA was extracted using a Total RNA Purification Kit (Norgen Biotek Corp, Canada), according to the manufacturer’s instructions. The quality of total RNA was measured using an RNA Screen Tape on an Agilent Tape Station 4150 (Agilent Technologies, USA). The quantity of total RNA was measured using a NanoDrop spectrophotometer (Thermo Fisher Scientific, USA) and a Qubit RNA High Sensitivity Assay on a Qubit 2.0 Fluorometer (Life Technologies, USA). The complete sequence library preparation and transcriptome sequencing for the Illumina NextSeq 500 protocol were performed by Novogene Co. Ltd. (Singapore). Raw RNA sequencing reads were deposited in the Sequence Read Archive (SRA) database of the National Center for Biotechnology Information (NCBI). All the sequences were listed in Table S2. After filtering, clean reads were mapped to the oil palm reference genome *E. guineensis* (GCF_ 00044 2705.1) using HISAT (v2.1.0) (Kim et al., 2015) with default parameters. StringTie (v1.3.4) (Pertea et al., 2015) was used to calculate read counts. The counts per million (CPM) values were used to quantify gene expression abundance and variation. DESeq2 (Love et al., 2014) was used to identify differentially expressed genes (DEGs) between the sample groups. Genes or transcripts with a false discovery rate (FDR) < 0.05 and |log2(fold change)| ≥ 2 were used as thresholds for significant differential expression. The Kyoto Encyclopedia of Genes and Genomes (KEGG) enrichment analysis was used to determine the pathways of significant enrichment in DEGs.

#### Native PAGE and Zymograms

A crude extract of mesocarp tissue for enzymatic activity was prepared as previously described (Malinova et al., 2014). Soluble starch synthase (SS) activity was measured as previously described (Brust et al., 2014) with minor modifications. The SS activity of samples at 22 WAP from the high (H), medium (M), and low (L) yields were measured in triplicate. Next, 50 µg of protein was loaded onto the native gel and run for 2 h at 4°C and 30 mA. The gel was incubated with 50 mM tricine-KOH (pH 8.0), 0.025% (w/v) bovine serum albumin, 5 mM dithioerythritol, 2 mM EDTA, and 25 mM potassium acetate. After 10 min, the incubation mixture was replaced with fresh incubation buffer, and 1 mM ADPglucose and 0.5% (w/v) soluble potato starch were added. Zymograms of SS activity were obtained by incubating the separation gels for 5 h at RT, followed by staining with Lugol solution (0.25% (w/v) I_2_ and 1% (w/v) KI).

### Validation of RNA-seq data using qRT–PCR

Total RNA was extracted from the mesocarp tissue using a total RNA purification kit (Norgen Biotek Corp, Canada). First-strand cDNA was synthesized using the Maxima First Strand cDNA Synthesis Kit (Thermo Fisher Scientific, Lithuania). All qRT–PCR reactions were carried out using SYBR Green PCR Master Mix (Applied Biosystems, USA) on an ABI 7500 Fast Real-Time PCR system (Applied Biosystems, USA) following the manufacturer’s instructions. Each reaction was performed in triplicate. The relative expression levels were determined using the 2^−ΔΔCt^ method, and β-actin was used as the reference gene. All the primers used are listed in Table S3.

## Supplemental data

These materials are available in the online version of this article.

Table S1. List of starch metabolism genes investigated in this study.

Table S2. List of NCBI database accession numbers and summary of sequencing reads after filtering.

Table S3. List of primers used for qRT–PCR validation.

Table S4. The list of differentially expressed genes (DEGs) during fruit development.

Table S5. The list of genes involved in glycolysis in oil palms.

Figure S1. KEGG enrichment analysis of DEGs in oil palm fruits at different developmental stages.

Figure S2. Heatmap analysis of starch metabolism-related genes during fruit development.

Figure S3. The detailed heatmap analysis of gene isoforms related to yield in the starch metabolism pathway.

Figure S4. Starch granule size distribution and differential chain length distribution at 22 WAP.

Figure S5. Activity staining of soluble starch synthase.

Figure S6. Heatmap analysis of glycolysis genes during fruit development.

Figure S7. Validation of RNA-seq results by qRT–PCR for selected genes.

## Acknowledgment

The authors thank the research and development team of PT. Astra Agro Lestari Tbk, Indonesia, for providing plant material and technical assistance. Special thanks to Reza Ernawan, M. Krisna A. Putra and Ricki Susilo for their kind help.

## Author contributions

AA and JF conceived of the project and designed the experiments. AA and JC conducted the experiments. AA analyzed the data and wrote the draft of the manuscript. JF supervised the experiments and revised the manuscript. All authors have read and agreed to the published version of the manuscript.

## Data availability statement

Raw sequencing reads used in this study have been deposited in the NCBI database under the Bioproject accession no. PRJNA882487. Reviewer link: https://dataview.ncbi.nlm.nih.gov/object/PRJNA882487?reviewer=r76pr591aqag1dt5pf4vmrdb28

